# Embracing enzyme promiscuity with activity-based compressed biosensing

**DOI:** 10.1101/2022.01.04.474983

**Authors:** Brandon Alexander Holt, Hong Seo Lim, Melanie Su, McKenzie Tuttle, Haley Liakakos, Peng Qiu, Gabriel A. Kwong

## Abstract

Genome-scale activity-based profiling of proteases requires identifying substrates that are specific to each individual protease. However, this process becomes increasingly difficult as the number of target proteases increases because most substrates are promiscuously cleaved by multiple proteases. We introduce a method – **S**ubstrate **Li**braries for **C**ompressed sensing of **E**nzymes (SLICE) – for selecting complementary sets of promiscuous substrates to compile libraries that classify complex protease samples (1) without requiring deconvolution of the compressed signals and (2) without the use of highly specific substrates. SLICE ranks substrate libraries according to two features: substrate orthogonality and protease coverage. To quantify these features, we design a compression score that was predictive of classification accuracy across 140 *in silico* libraries (Pearson *r* = 0.71) and 55 *in vitro* libraries (Pearson *r* = 0.55) of protease substrates. We demonstrate that a library comprising only two protease substrates selected with SLICE can accurately classify twenty complex mixtures of 11 enzymes with perfect accuracy. We envision that SLICE will enable the selection of peptide libraries that capture information from hundreds of enzymes while using fewer substrates for applications such as the design of activity-based sensors for imaging and diagnostics.

## Introduction

Proteases are a major class of enzymes; more than 600 enzymes, comprising ~3% of the human genome^1^, are classified as proteases due to their ability to hydrolyze peptide bonds and degrade proteins (i.e., proteolysis). Protease activity is a driver of important biological processes, ranging from development and differentiation^2^ to pathological conditions such as cancer, neurodegenerative disorders, and inflammatory diseases^3^. However, due to the irreversible nature of proteolysis, protease activity is tightly regulated via mechanisms such as inhibitory prodomains, cofactor binding, and protein inhibitors^4^. Given this degree of posttranslational regulation, quantifying protease activity, rather than transcriptomic or proteomic analyses, is often required to understand the biological roles of proteases^5^. This has motivated the development of activity-based sensors – probes that quantify protease activity – which are used for early detection of disease^6–11^, biological imaging^12–14^, and drug screening^15, 16^. The two primary compositions of activity-based sensors are (1) substrates that produce a signal upon proteolysis and (2) probes that bind active proteases^17^. For the former approach, a major bottleneck is substrate design, which involves screening for peptide substrates that are specific to the target protease (**Fig. 1**, step 1). However, finding substrates with high specificity becomes increasingly difficult as the number of target enzymes increases because most proteases are characterized by promiscuous activity^18–26^.

**Figure 1.**
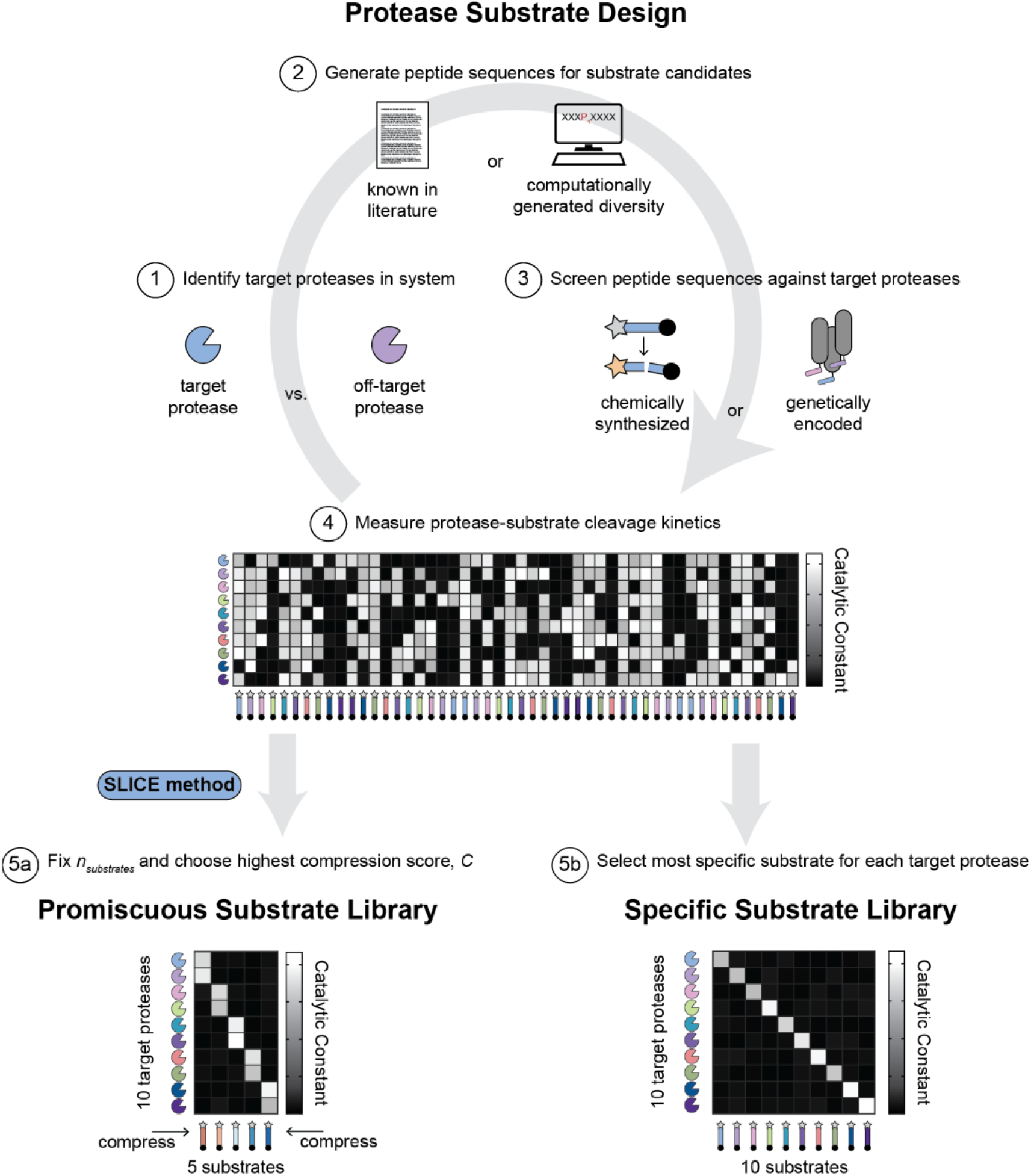
Conceptual overview of protease substrate design using the SLICE method. (1) Identify which proteases in the system being probed are considered target proteases (blue pac man), and which are off-target proteases (purple pac man). (2) Generate candidate peptide sequences that can be used as substrates for target proteases. Peptide sequences can be acquired from the literature (paper icon) or computationally generated (computer icon). Computationally generated diversity includes degenerate libraries as well as predicted sequences derived from computational modeling software. (3) Screen candidate peptide sequences against all protease targets via chemically synthesized activity-based sensors (e.g., fluorogenic probes, peptide microarrays, etc.) or genetically encoded libraries (e.g., phage display, bacteria display, etc.). (4) Heat map of cleavage kinetics, quantified by the catalytic constant, *k_cat_*, for all protease-substrate pairs (rows = proteases, columns = substrates). (5a) An example promiscuous substrate library which has fewer substrates (*n_substrates_* = 5) than proteases (*n_proteases_* = 10). The compression score, *C*, represents the score assigned to the library by the SLICE method, with 1 being the highest score and 0 the lowest. (5b) An example specific substrate library which has the same number of substrates as proteases (*n_substrates_* = *n_proteases_* = 10).

To accelerate the process of designing specific substrates, methods to generate and screen libraries of peptide sequences have been developed, including positional scanning libraries^27–29^, peptide microarrays^30, 31^, fluorogenic peptides^32, 33^, and other mixture-based peptide libraries^34, 35^. These libraries are either degenerate or diversified at certain positions based on consensus cleavage motifs from the literature^36^ or computational approaches to predict peptide sequences based on the structure of the active site of a target protease^37, 38^ (**Fig. 1**, step 2). To generate potentially novel specific substrates, high-throughput evolution-based methods display and iteratively screen randomized peptide sequences on the surface of bacteria (e.g., CLiPS)^39, 40^ or bacteriophages (e.g., phage display)^41^, and have been extended for screening endogenous protease activity*^42, 43^* (**Fig. 1**, step 3—4). To further increase substrate specificity, approaches have been developed to broaden the chemical diversity of peptide libraries, such as via the introduction of non-natural amino acids^44, 45^ or cyclic peptide libraries^46–49^. In cases where protease-substrate kinetics are known, signal deconvolution algorithms can retroactively infer the activity levels of individual enzymes in a complex mixture^9, 32^; this approach works well on controlled reactions involving recombinant enzymes. With these methods, libraries of up to 10—20 substrates, each of which have unique molecular barcodes, have been constructed to sense dysregulated protease activity for early detection of disease^50–53^. However, the current paradigm in substrate design methods is to favor specific substrates over promiscuous candidates.

Here, we embrace enzyme-substrate promiscuity by developing a substrate design method – **S**ubstrate **Li**braries for **C**ompressed sensing of **E**nzymes (SLICE) – for selecting complementary promiscuous substrates to compile libraries of activity-based sensors that can classify distinct protease mixtures without specific substrates or signal deconvolution (**Fig. 1**, step 5). Rather, SLICE, inspired by the signal processing technique compressed sensing^54–57^, evaluates different combinations of substrates to find the most complementary library that maximally senses all target proteases. We accomplish this by designing a compression score, *C*, which scores substrate libraries according to two features: (1) substrate orthogonality, which measures the uniqueness of protease-substrate kinetics and (2) protease coverage, which measures the total fraction of target proteases sampled. In a simulated disease detection challenge based on a melanoma gene microarray dataset^58^, the compression score was predictive of classification accuracy across 140 *in silico* libraries (Pearson *r* = 0.71) and 55 *in vitro* libraries Pearson *r* = 0.55). Further, we used SLICE to design a 2-substrate library (*C* = 0.94) that classified 20 complex samples containing one of two distinct 11-protease mixtures *in vitro* with perfect accuracy. Looking forward, producing smaller libraries will reduce the number of read-outs, overall cost, and processing time, which is ideal for imaging and activity-based diagnostics. We envision that SLICE will enable the selection of promiscuous substrate libraries that capture information from hundreds of enzymes using fewer activity-based sensors than is currently possible.

## Results

### Computational pipeline for evaluating classification performance of simulated substrate libraries

Given an initial pool of candidate substrates, our goal was to develop a method for predicting which libraries of promiscuous substrates should be selected to accurately classify distinct mixtures of proteases. Therefore, we sought to create a simulation pipeline for evaluating the classification performance of substrate libraries with known protease-substrate cleavage kinetics (e.g., catalytic constants, *k_cat_*). To simulate a disease detection problem, we used a microarray gene expression dataset^58^ for 162 extracellular proteases in a murine melanoma model, (**Fig. S1a**) and generated Gaussian-distributed populations of 200 simulated samples from a healthy condition (Day 1) and a disease condition (Day 7) (i.e., 100 simulated samples for each condition) (**Fig. 2**, part 1a). After performing principal component analysis on the simulated samples, we observed that the first two principal components represent >80% of variance and provide a clear separation between the healthy and disease groups, meaning that the two groups can be easily classified using all protease measurements simultaneously. Given the challenge of sensing the activity of all proteases simultaneously, we use libraries of promiscuous substrates to measure combinations of proteases. To simulate promiscuous substrate libraries, we randomly generated catalytic constants, *k_cat_*, for all pairwise combinations of proteases and substrates (**Fig. 2**, part 1b). We calculated the product formation rates, *V_max_*, for each substrate across all simulated samples by multiplying the protease gene expression, *P*, by the catalytic constant, *k_cat_*, for each protease-substrate pair (**Fig. 2**, part 2). We used a random forest model or classifying the simulated samples (i.e., healthy vs. disease) and used 5-fold cross-validation by aggregating predictions of an unseen fold (i.e., test set) based on the model trained by the other four folds (i.e., training set). To quantify classification performance, we calculated the area under the receiver operating characteristic curve (AUROC) resulting from applying the trained model to the test set (**Fig. 2**, part 3; **Fig. S1b**). We observed a clear trend that increasing the number of substrates in a library resulted in increased classification power. With this pipeline, we can evaluate the classification performance for a substrate library with known catalytic constants in a simulated disease detection problem as a proxy for true classification power.

**Figure 2.**
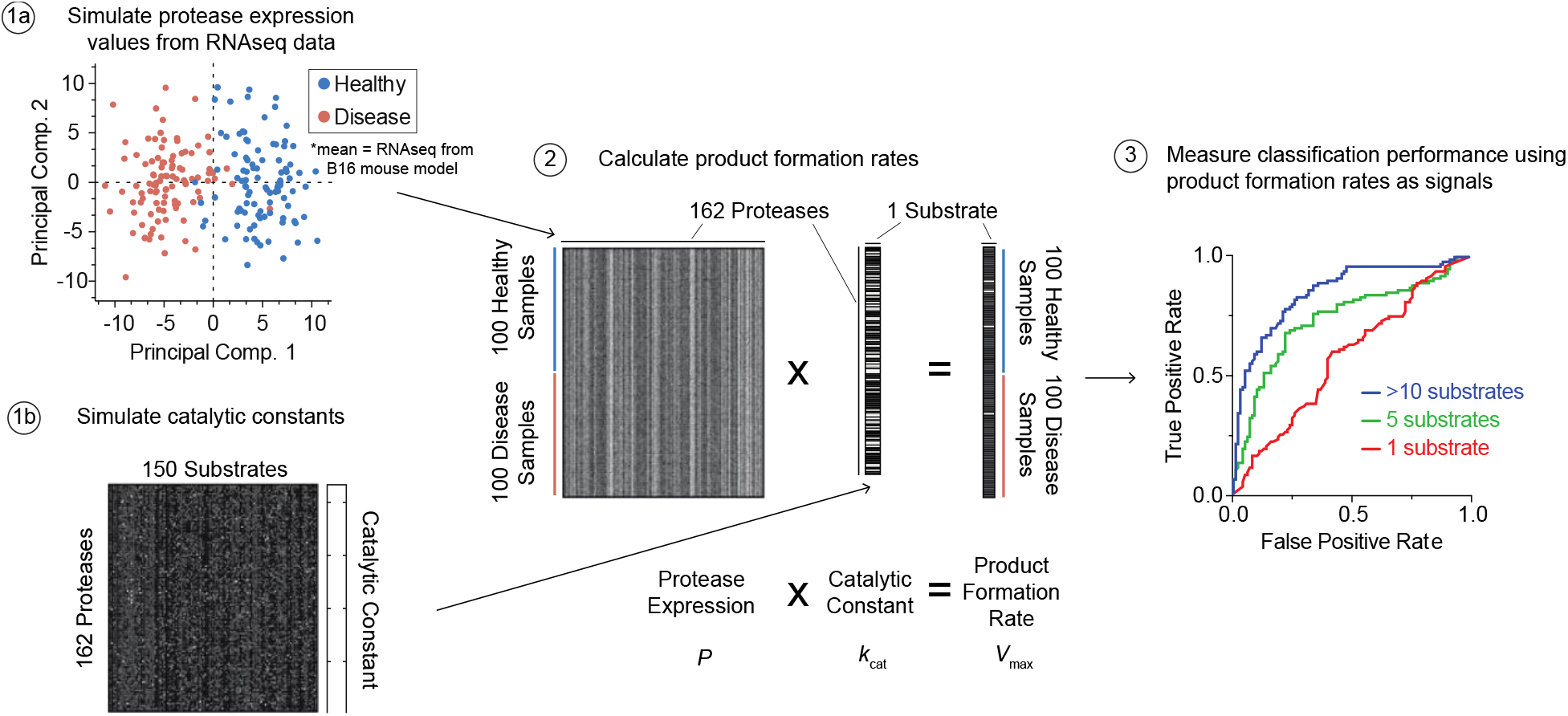
Computational pipeline for evaluating classification performance of simulated substrate libraries. (1a) Plot of first two principal components from principal component analysis on microarray gene expression data of 162 protease genes in Day 1 (healthy, blue) and Day 7 (disease, red) mouse tissue samples in a B16 melanoma model. To simulate 100 samples and 100 disease samples are computationally generated as a gaussian distribution from a single biological sample. (1b) Heat map of simulated catalytic constants, *k_cat_*, for every pairwise combination between 162 proteases and 150 substrates (white = high, black = low). (2) Visualization of how product formation rates, *V_max_*, are calculated using protease concentrations, *P*, and catalytic constants, *k_cat._* The result of this calculation is a product formation rate per substrate per sample. (3) Receiver operating characteristic (ROC) curves as a measure of healthy vs. disease classification performance using product formation rates as features of observations used to train a random forest model. Blue trace is ROC curve when using signals (i.e., product formation rates) from 11 substrates (green trace = 5 substrates, red trace = 1 substrate).

### A compression score for promiscuous substrate library selection

Since a promiscuous substrate can be cleaved by multiple proteases, the net signal of a substrate represents some weighted combination of product formation rates from multiple proteases. Therefore, measuring the signal of a promiscuous substrate compresses the product formation rates (i.e., activity) from multiple proteases into one feature. We sought to design a compression score, *C*, that selects for the most complementary set of promiscuous substrates that maximally senses the proteases-of-interest. To account for this, the compression score is a weighted sum of two metrics - substrate orthogonality, *S_orth._*, and protease coverage, *P_cov._* (**Fig. 3a, Equation 1**).

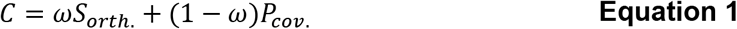

**Figure 3.**
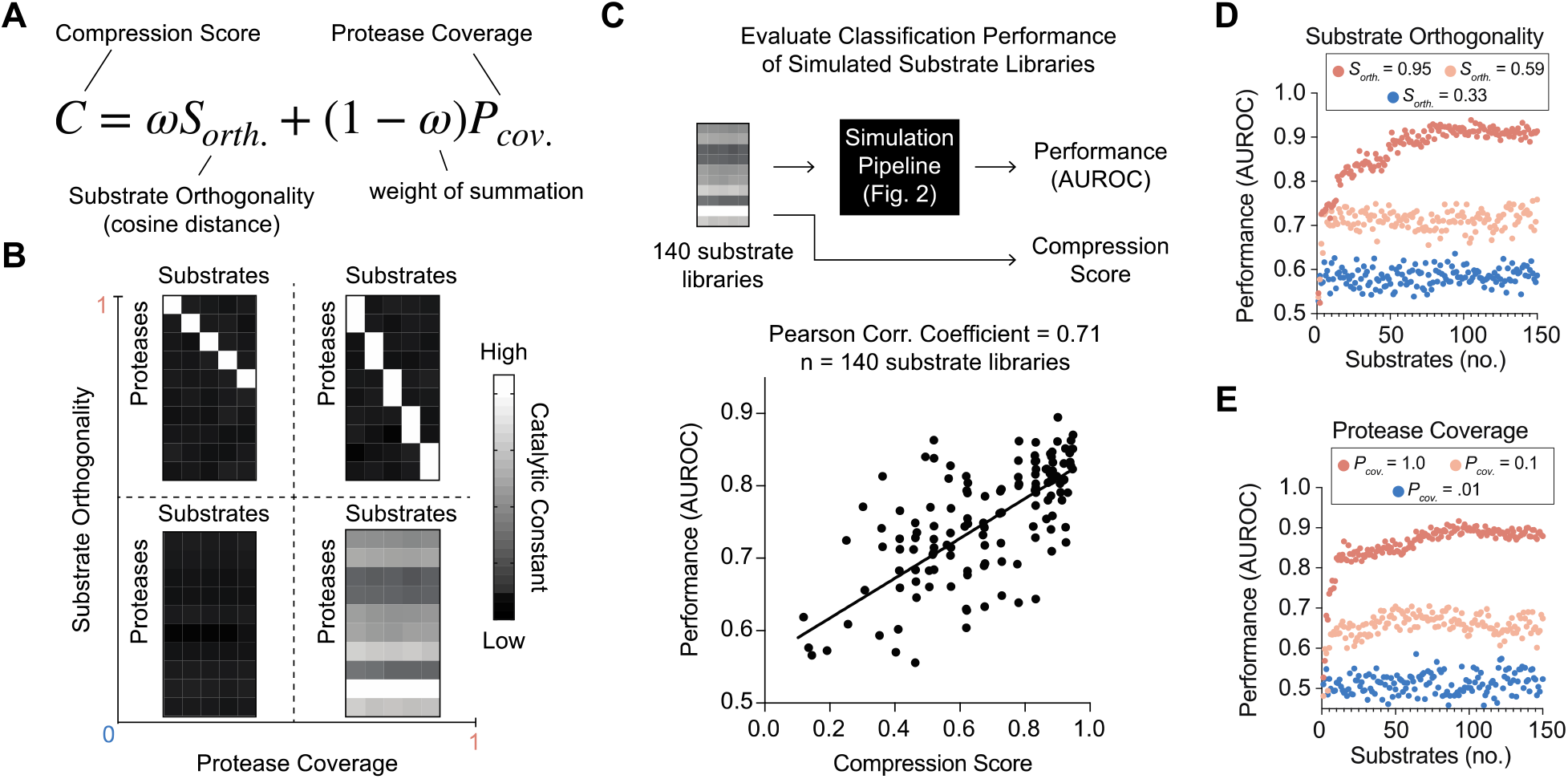
A compression score for promiscuous substrate selection. **(A)** Equation used to calculate the compression score, *C*. Substrate orthogonality, *S_orth._*, which is quantified by the cosine distance metric, and protease coverage, *P_cov._*, which quantifies the fraction of proteases that are sampled by a substrate library, are combined according to the weight of summation, ω. All variables range from 0 to 1. **(B)** Schematic showing four example substrate libraries and their relative magnitude in substrate orthogonality (y-axis) and protease coverage (x-axis) space. Each substrate library is represented with a heat map of catalytic constants (white = high, black = low) for all protease (rows) and substrate (columns) combinations. **(C)** (top) Schematic showing pipeline for calculating the compression score and classification performance for 140 simulated substrate libraries. (bottom) Plot of correlation between the compression score (x-axis) and classification performance (AUROC, y-axis). Black line is line of best fit. Each dot represents the performance of one substrate library averaged over 5 repeats. Plots showing classification performance (AUROC, y-axis) vs. substrate library size (number of substrates, x-axis) for changing value of substrate orthogonality **(D)** and protease coverage **(E).** Each dot represents the performance of one substrate library.

The compression score operates on a 2D matrix of kinetic constants (e.g., catalytic constants, product formation rates, etc.) for all pairwise combinations of protease (rows) and substrate (columns); the score outputs one value ranging between 0 and 1, with 1 being the optimal score (**Fig. 3b**). Substrate orthogonality, *S_orth._*, which is the cosine distance metric (**Fig. S2**), quantifies the orthogonality of the columns, or how unique each of the substrates are from one another in the protease space. For example, substrate libraries with high *S_orth._* will have columns that are different from one another, whereas the columns will be more similar in libraries with low *S_orth._* (**Fig. 3b**, y-axis). Conversely, protease coverage quantifies how many rows have at least one element with a high value, or how many proteases are collectively sampled by a library. For example, substrate libraries with high *P_cov._* will have a high value in all rows, whereas libraries with low *P_cov._* will include rows of only low values (**Fig. 3b**, x-axis). To verify that the compression score is predictive of classification performance, we used the computational pipeline described in Figure 2 to evaluate the classification performance of 140 simulated substrate libraries. We found that the compression score demonstrated a strong correlation with classification performance (Pearson’s r = 0.71), with substrate libraries where *C* < 0.2 provided little useful information (i.e., 0.5 < *AUROC* < 0.6) and libraries where *C* > 0.9 demonstrated strong classification performance (i.e., *AUROC* > 0.85) (**Fig. 3c**). To verify that both *S_orth._* and *P_cov._* contribute to the compression score independently, we independently fixed each variable and observed the change in classification performance across varying substrate library sizes (i.e., 1 < *n_substrates_* < 150). We found that increasing both substrate orthogonality (**Fig. 3d**) and protease coverage (**Fig. 3e**) independently increased classification performance from 0.5—0.6 to >0.9 across all substrate library sizes tested. With the compression score, we can rank-order and select the optimal set of promiscuous substrates where the kinetic constants towards the relevant protease targets are known.

### Exhaustive scoring of substrate libraries *in vitro* with SLICE

To demonstrate the process of constructing a substrate library with SLICE experimentally, we selected 11 target proteases, comprising primarily metallo- and serine proteases, and a candidate pool of 11 substrates, with known cleavage activity from matrix metalloproteases (MMP) and cathepsin proteases (**Table S1**). We designed fluorogenic probes for these substrate sequences by flanking each with a fluorophore and quencher such that peptide cleavage would result in a measurable increase in fluorescence (**Fig. 4a**, part 1). We performed cleavage assays for all 121 unique protease-substrate pairs and extracted the product formation rates as representative kinetic parameters (**Fig. 4a**, part 2; **Fig. S3**). We observed that although the substrate sequences were known to target MMPs and cathepsins, the off-target proteases (e.g., KLK2, thrombin, etc.) used in these experiments also showed the propensity to cleave these sequences, which can be attributed to the promiscuous nature of protease substrates. To visualize the distribution of scores for these libraries, we exhaustively enumerated all libraries with sizes ranging from 2 to 10, and computed the *S_orth._, P_cov._*, and *C* scores for all those libraries (**Fig. 4b**, part 1). We found that this candidate pool of substrates produced libraries high in *P_cov._* but low in *S_orth._* (**Fig. 4b**, part 2). We found that the mean *C* score of 0.66 (n = 2,035 libraries) was higher than the benchmark score of randomly-generated libraries (*C* = 0.6), meaning that real substrates tended to be more promiscuous than randomly-generated substrates (**Fig. 4b**, part 3). To validate that the compression score is predictive of substrate library performance using empirically derived kinetic constants (i.e., product formation rates), we repeated the pipeline described in **Fig 2** using the product formation rates found in **Fig. 4a**. We trimmed down the list from 162 to 11 protease genes that were either from the same family or an exact match to the 11 proteases used in our experiments and simulated 100 healthy and 100 disease samples (**Fig. 4c**, part 1). To fix library size, we calculated the distribution of compression scores of all libraries comprising only two substrates (**Fig. 4c**, part 2), and found that this distribution closely matched the score distribution for all library sizes (**Fig. 4b**, part 3). We evaluated the classification performance for all 55 libraries of size 2 and found that the compression score correlated with AUROC (**Fig. 4c**, part 3; Pearson’s *r* = 0.55). Here, we demonstrated that constructing a library with SLICE involves (1) selecting a candidate pool of substrates that broadly recognize known protease targets (2) measuring a kinetic parameter for each protease-substrate pair, and (3) identifying the optimal library/libraries by evaluating the compression score.

**Figure 4.**
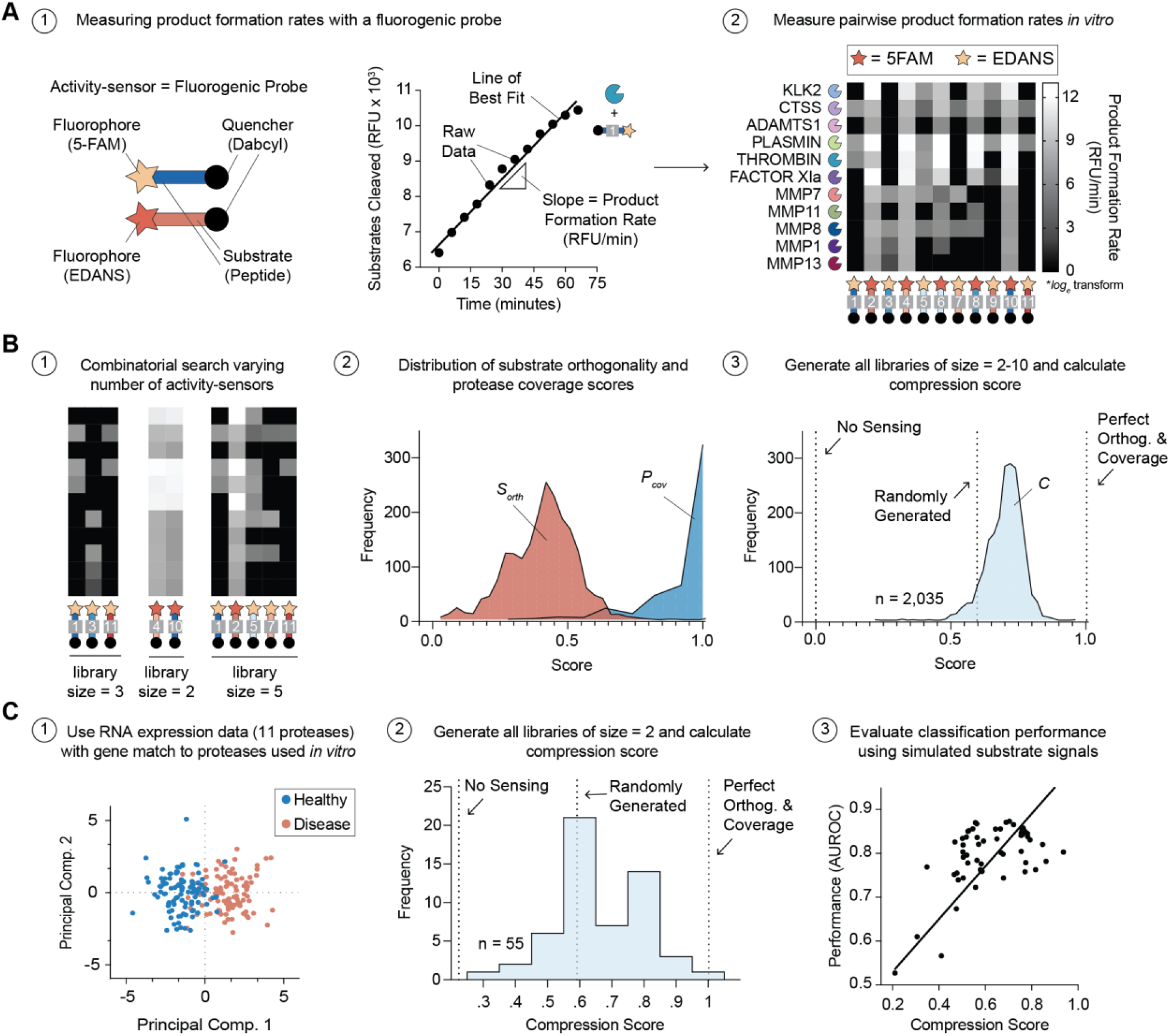
Exhaustive scoring of substrate libraries *in vitro* with SLICE. **(A)** (1, left) Schematic of activity-sensor or fluorogenic probe. Activity-sensor comprises a peptide substrate (blue and red bar) flanked with a fluorophore (yellow star = 5-FAM, red star = EDANS) and a quencher (black circle = Dabcyl). Upon cleavage, the fluorophore and quencher are separating, which results in an increase in fluorescent signal. (1, right) Cleavage assay of thrombin and substrate-1 showing the increase in number of substrates cleaved (y-axis) over time (x-axis). Black dots are raw data. The slope (triangle) of the line of best fit (black line) is calculated as the product formation rate. Units of RFU/min are used, as RFU correlates with the number of substrates cleaved. (2) Heat map showing all pairwise combinations of product formation rates as measured from independent cleavage assays. Proteases are in rows and substrates are in columns. Data is natural log transformed. **(B)** (1) Schematic showing that all unique combinations of substrates, with library size ranging from 2 to 10, are scored with SLICE. (2) Histogram showing the distribution of substrate orthogonality (*S_orth_*, red distribution) and protease coverage (*P_cov_*, blue distribution) scores. (3) Histogram showing the distribution of compression scores (*C*, light blue distribution). Vertical dashed lines depict the score of various controls. ‘No Sensing’ depicts the score of a library where kinetic constant = 0 for all protease-substrate pairs. ‘Randomly Generated’ depicts the score of a library where kinetic constants are randomly generated. ‘Perfect Orthog. & Coverage’ depicts the score of a library where all proteases are sampled, and each substrate has no overlapping kinetic constants. **(C)** (1) Principal component analysis of 11 proteases selected from 162 found in original B16 study. Proteases selected as either exact match or as member of same family as 11 proteases used in our study (A, part 2). Each dot represents one simulated sample (red = disease, blue = healthy). (2) Histogram showing the distribution of compression scores (light blue distribution) for all substrate libraries of size 2 (i.e., 2 substrates). (3) Plot showing correlation between compression score (x-axis) and classification performance (y-axis, AUROC). Black line shows line of best fit.

### Experimental demonstration and validation of substrate library design with SLICE

To validate the efficacy of a promiscuous substrate library designed with SLICE, we created an *in vitro* classification challenge for detecting dysregulated protease activity. To represent the two classification groups (i.e., protease mixture A vs. protease mixture B; **Fig. 5a**, part 1), we randomly generated two distinct mixtures of the same 11 target proteases from previous experiments (**Fig. 3, 4**). We incubated the library separately with 10 hand-pipetted repeats of both mixtures to introduce variance in the protease concentrations within the same group (**Fig. 5a**, part 2). To evaluate the classification performance, we used the product formation rates of each substrate as the observations used to train a random forest model and calculated the AUROC for all test set samples in all 5-fold cross-validation iterations (**Fig. 5a**, part 3). As a negative control, we tested a library with a low compression score (*C* < .5) to benchmark the performance of the SLICE library (*C* > .9) (**Fig. 5a**). The kinetic parameter heatmap for the SLICE library (*C* = .95) showed that there is at least one substrate that can sense each protease, and the substrates only overlapped on one protease target (i.e., MMP8). Conversely, the negative control library (*C* = .49) does not sense 3 proteases (i.e., MMP1, MMP7, MMP13) and the substrates overlap on 4 protease targets (i.e., KLK2, CTSS, Plasmin, Factor XIa) (**Fig. 5b**). These results validate that the scoring system (i.e., compression score, *C*) used in the SLICE method accurately represents protease coverage and substrate orthogonality (**Fig. S4**). After incubating each library with all 20 protease mixtures (i.e., 10 repeats of mixture A, 10 repeats of mixture B), we plotted the results from each mixture in substrate space (i.e., x-axis = product formation rate of 5-FAM substrate, y-axis = EDANS substrate) (**Fig. S5**). We observed that the SLICE library (*C* = .95) provided strong separation between mixture A and mixture B when compared to the negative control (*C* = .49) (**Fig. 5c**). These results were confirmed by AUROC analysis, where the SLICE library (*C* = .95) classified all twenty mixtures with perfect accuracy (AUROC = 1.00), while compressing the dimensionality from 11 proteases to 2 substrates. By comparison, the negative control library (*C* = .49) showed worse classification performance (AUROC = .58), which held true across all temporal endpoints tested (**Fig. 5d; Fig. S6**). Further, we found that the same substrate signal (i.e., substrate-8) that resulted in a negative feature importance score in the negative control (*C* = .49) library, produced a positive feature importance score in the SLICE (*C* = .95) library (**Fig. S7**). This demonstrates that while promiscuous substrates can be detrimental to certain libraries, pairing them with complementary substrates can improve the overall classification performance of the library. Here, we demonstrated that the SLICE method can select for substrate libraries and assign a compression score that accurately predicts their classification performance when differentiating complex protease activity.

**Figure 5.**
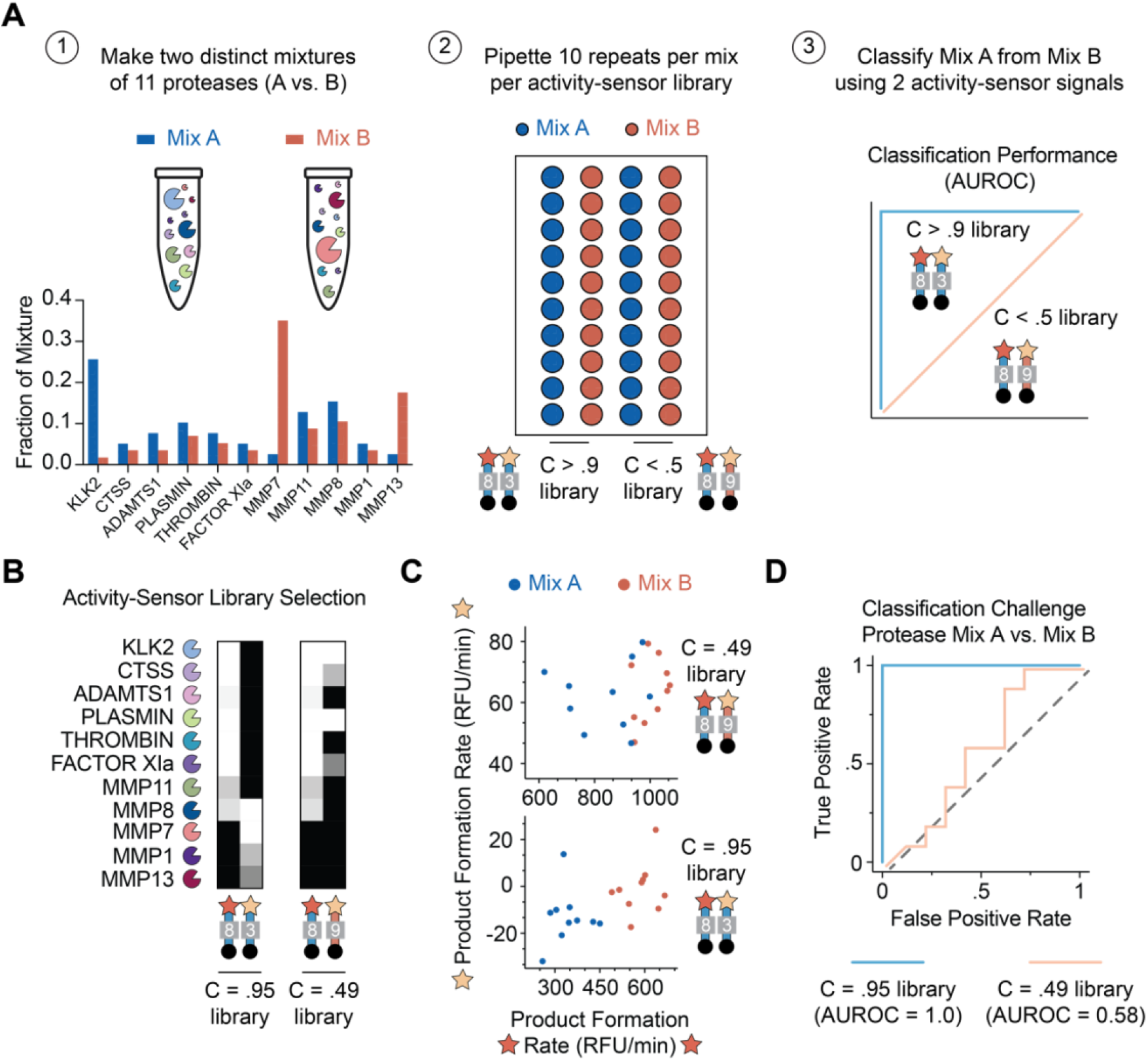
Experimental validation of substrate library design with SLICE. **(A)** Schematic of experimental workflow: (1) Two mixtures (A = blue, B = red) of 11 proteases are randomly generated. Each mixture is represented with a test tube containing 11 proteases (pac man shape). Relative size of protease roughly represents the relative concentration. Actual relative concentrations are plotted in bar graph below (A = blue bars, B = red bars). (2) Schematic of experimental well-plate containing samples of protease mixtures (1 circle = 1 well). Both mixtures are independently pipetted 10 times each (blue well = Mix A, red well = Mix B) to create a population with variance due to pipetting error. One library is introduced to all 20 samples (10 of mixture A, 10 of mixture B) and the product formation rates of both activity-based sensors in the library are measured. (3) Schematic graph (not real data) showing that the library with a high compression score (C > .9) should have high classification performance (blue line) whereas the library with low compression score (C < .5) should have low classification performance (orange line). **(B)** Heat maps showing the product formation rates for the library with the highest compression score (C = .95 library) and the library with the lowest compression score (C = 0.49 library). (white = high product formation rate, black = low product formation rate). **(C)** Plot of the resulting product formation rates for each activity-sensor after incubation with protease mixtures (1 dot = 1 mixture; blue dot = mixture A, red dot = mixture B). The product formation rates from activity-based sensors using 5-FAM are plotted on the x-axis, and product formation rates from EDANS are plotted on the y-axis. The top plot shows the results when using the C = .49 library, and the bottom plot shows the results when using the C = .95 library. **(D)** AUROC plot showing the results of classifying mixture A from mixture B when using the C = .95 library (blue trace) or the C = .49 library (orange trace).

## Discussion

Here, we develop a method, SLICE, for compiling libraries of promiscuous substrates that sense protease activity for classification or diagnostic applications. This method involves (1) selecting a candidate pool of substrates that sense the target proteases, (2) measuring a kinetic parameter (e.g., *k_cat_, V_max_*, etc.) for each protease-substrate pair, and (3) identifying the optimal library of a fixed size by evaluating the compression score. The advantages of this method are that it enables the use of fewer promiscuous substrates (i.e., specific substrates not required) than the number of target proteases. By comparison, the current paradigm is to search for substrates that are specific to one protease^8, 59^ and use approximately the same number of substrates as proteases^50^. With these methods, all off-target protease activity is considered background noise, which is traditionally filtered out via chemical^46, 48^ or computational methods^9, 32^. As the number of enzyme targets increases, it becomes increasingly difficult to maintain specificity across all substrates. Further, since each substrate requires a unique reporter, the number of simultaneous read-outs becomes limited by cost (e.g., mass barcodes^50^) or physical restrictions (e.g., fluorescence^15^).

It is suggested that protease promiscuity bolsters fitness by (1) providing alternative evolutionary starting points and (2) increasing biological efficiency (i.e., multiple functions per enzyme)^26^. We proposed that by embracing protease promiscuity could leverage the ubiquity of substrates that recognize multiple targets. Serving as inspiration for the SLICE method, compressed sensing (CS) is a signal processing technique that utilizes measurements of a mixture of multiple target signals to recover information of individual signals^54^. A famous application of CS is the single-pixel camera, which demonstrated the ability to efficiently handle high dimensional datasets (e.g., hyperspectral imaging, video, etc.) and inspired the use of CS in magnetic resonance imaging^55^ and imaging transcriptomics^56, 57^. CS utilizes compressed signals, which are a composite of multiple different signals; this mirrors how the total number of cleaved copies of a promiscuous substrate results from a weighted combination of different proteases. However, a major difference is that our method does not require the deconvolution of compressed signals (i.e., cleaved substrate signals). Future iterations of SLICE could incorporate (1) CS features (e.g., sparsity, incoherence) for substrate selection metrics (i.e., compression score) and (2) deconvolution of the compressed signals. However, we found that compressed signals are often sufficient for achieving high classification accuracy and would be preferable for applications such as point-of-care^60^ or imaging diagnostics^57^ where fewer signals reduces overall cost and processing time.

We envision that SLICE will be a useful for applications where obtaining precise activity values per protease is less important than detecting systems-level changes, such as disease-staging, classification, and diagnosis. The ultimate application of SLICE would be a universal substrate library that is constructed by running all candidate substrates though a standardized test, which measures *k_cat_* against all >600 recombinant human proteases. From this library, various sub-libraries targeting different groups of proteases could be extracted on a per-application basis. For example, a diagnostic activity-sensor library could be extracted from the universal library by defining disease-specific target proteases ideally in pathologies that can be diagnosed using blood or plasma samples, such as coagulation disorders^61, 62^ or cancer^63, 64^. While *in vitro* protease activity measurements may not fully account for the dynamic states of proteases *in vivo^65,66^*, future work could improve this by creating more robust *in vitro* tests that sample proteases under multiple states (e.g., redox, fluid dynamics, etc.) or developing *in vivo* tests that isolate the activity form individual proteases.

Further, other classes of enzymes also exhibit promiscuity^67^, which means the design rules presented in this work can likely be extended to other promiscuous enzymes such as kinases or phosphatases and their activity-based sensors^68–73^. For example, candidate substrates would be mapped onto sensors that exhibit phosphorylation-or dephosphorylation-dependent changes in signal (e.g., fluorescence)^69, 74^. These sensors would be used to measure enzyme-substrate kinetics and generate an activity matrix, which could be processed using the SLICE method. In conclusion, we present SLICE as a method for embracing the use promiscuous substrates for detecting changes in protease activity, as an alternative approach to the use of specific substrates. Given the ubiquity of promiscuous substrates and the motivation to sense biological activity, we anticipate that the ideas presented here will have broad applicability to the field of enzyme-sensing at large.

## Acknowledgements

This work was funded by an NIH Director’s New Innovator Award (Award No. DP2HD091793), an R01 from the NCI (GR10003709), a U01 from the NCI and NIBIB (1U01CA265711), as well as projects from the NSF (CCF1552784 and CCF2007029). B.A.H is supported by the NSF GRFP (Grant No. DGE-1650044), National Institutes of Health GT BioMAT Training Grant under Award Number 5T32EB006343 and the Georgia Tech President’s Fellowship. G.A.K. holds a Career Award at the Scientific Interface from the Burroughs Welcome Fund. P.Q. is an ISAC Marylou Ingram Scholar, a Carol Ann and David D. Flanagan Faculty Fellow, and a Wallace H. Coulter Distinguished Faculty Fellow. The content is solely the responsibility of the authors and does not necessarily represent the official views of the National Institutes of Health. The authors thank Dr. Melissa Kemp (Georgia Tech & Emory) for their helpful discussions regarding the manuscript.

## Data Availability

The authors declare that the data supporting the findings of this study are available within the paper and its supplementary information files.

## Code Availability

The authors declare that the code supporting the findings of this study is available at https://github.com/brandon-holt/compressed-biosensing

## Supplementary Information

### Methods

#### Cleavage Assays

All protease cleavage assays were performed with a BioTek Cytation 5 Imaging Plate Reader, taking fluorescent measurements at 485/528 nm (excitation/emission) for read-outs measuring peptide substrates terminated with 5FAM (5-Carboxyfluorescein). In all conditions, substrate (20 μM) was added to protease (250 nM) in PBS for each well of a 384-well microplate immediately before reading began. Kinetic measurements were taken every minute over the course of 60 – 120 min at 37 C. Activity RFU measurements were normalized to time 0 measurement, and as such later time points (after time-0) represent fold change in signal. All fluorogenic peptide substrates were purchased from Genscript.

#### Simulation pipeline for evaluating classification performance of simulated libraries

Mouse protease expression data across different days (day 1, day 3, day 5, day 7) are used as the basis of the original simulation. Data from day 1 represents protease expression of relatively ‘healthy’ state, while data from day 7 represents protease expression of more severe tumor stage, hence representing ‘disease’ state. There are a total of 162 proteases measured across the days. The protease expression data from day 1 and day 7 are used to generate simulated data. 100 datapoints are randomly generated where each data is simulated by adding the random gaussian noise centered around 0 with a standard deviation of 2 to the protease expression data. We generated 100 healthy samples based on day 1, and another 100 disease samples based on Day 7, a total of 200. The subsequent dimension of each dataset is 162 by 100. Upon the generation of simulated datasets, we simulated substrates. For each simulated substrate, the number of the protease the substrate will sense is randomly sampled between 10 to 30, and the set of proteases is randomly chosen among the 162 proteases. We created a vector with a length of 162, where each element(protease) is assigned 0 if not chosen, and 1 if chosen. We simulate signals from the substrate probe by taking matrix multiplication of a transpose of the simulated protease data (100*162) and the substrate vector (162*1) resulting in a single signal value per sample. Hence, per one simulated substrate, we obtained 200 values where 100 belonging to healthy group and another 100 belonging to disease group. We created 150 such simulated substrates resulting in a total 150 signals per sample (150*200). The final signal data is used as an input to Random Forest (MATLAB TreeBagger; https://www.mathworks.com/help/stats/treebagger.html) to quantify whether samples could be classified as healthy or disease based on the signal data from the simulated substrates. 5-fold cross validation is applied to the dataset, and each fold left out during the training is applied to the model to evaluate probabilities belonging to either healthy or disease. The probability measures for the left-out folds are pooled together, and its accuracy and the area under the receiver operating characteristic (AUROC) are evaluated.

**Table S1.**
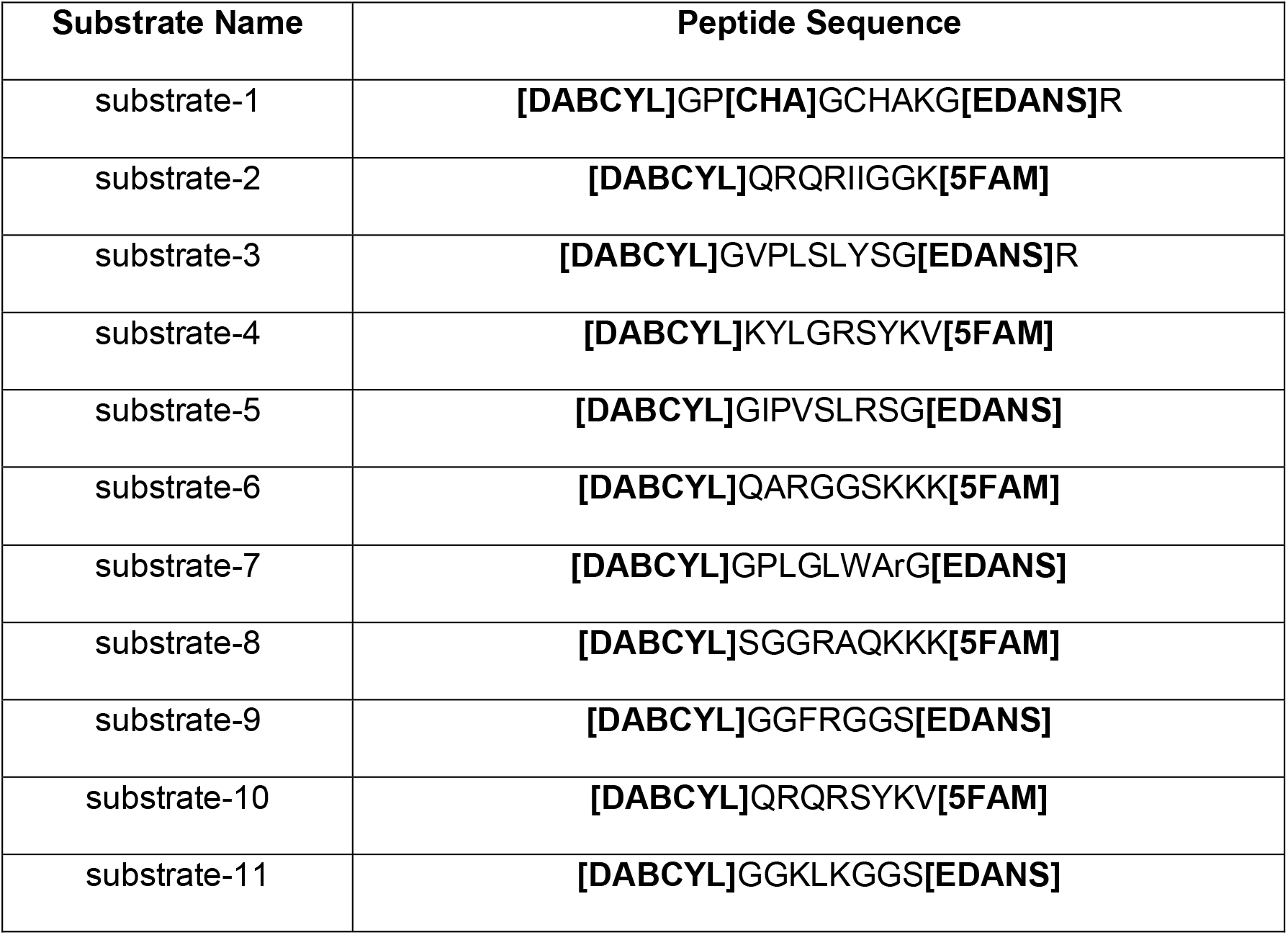
Substrate peptide sequences. Non-bold, capital letters represent single-letter L-amino acid codes. Lower case letters are D-amino acid codes. Bold letters represent in brackets functional groups: **[DABCYL]** = 4-(dimethylaminoazo)benzene-4-carboxylic acid **[EDANS]** = (5-((2-Aminoethyl)amino)naphthalene-1-sulfonic acid) **[5FAM]** = 5-Carboxyfluorescein **[CHA]** = (S)-2-Amino-3-cyclohexylpropanoic acid

**Figure S1.**
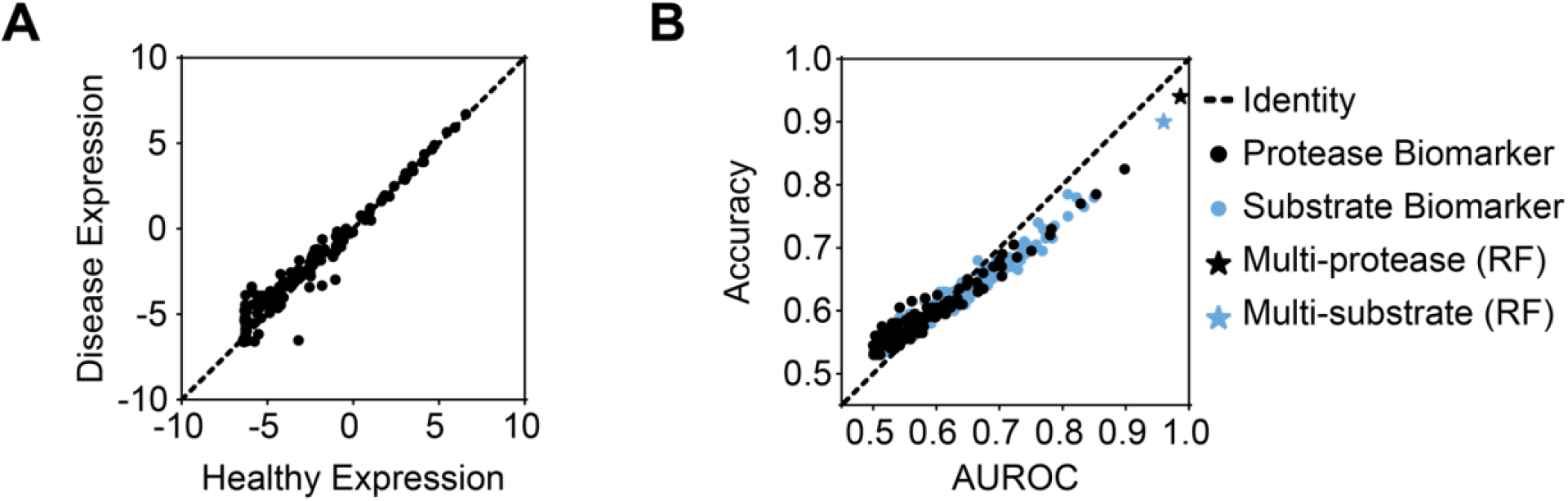
Plotting raw protease expression data and comparing healthy vs. disease classifiers. **(A)** Plot of healthy expression (x-axis; day 1) vs. disease expression (y-axis; day 7) of microarray gene expression data from B16 model found in literature. Each dot represents one protease gene, with a total of 162 extracellular protease genes. **(B)** Comparing various classifiers of healthy vs. disease expression profile in simulated population based on panel A. Area under the receiver operating characteristic (x-axis; AUROC), which results from a random forest model trained to classify healthy vs. disease, is compared to classification accuracy (y-axis). Blue and black dots represent the classification power of using the signal from single substrates or proteases, respectively. Blue and black stars represent the best multi-substrate or multi-protease classifiers, respectively. The dashed line is the identity line, where accuracy and AUROC are equal at all points.

**Figure S2.**
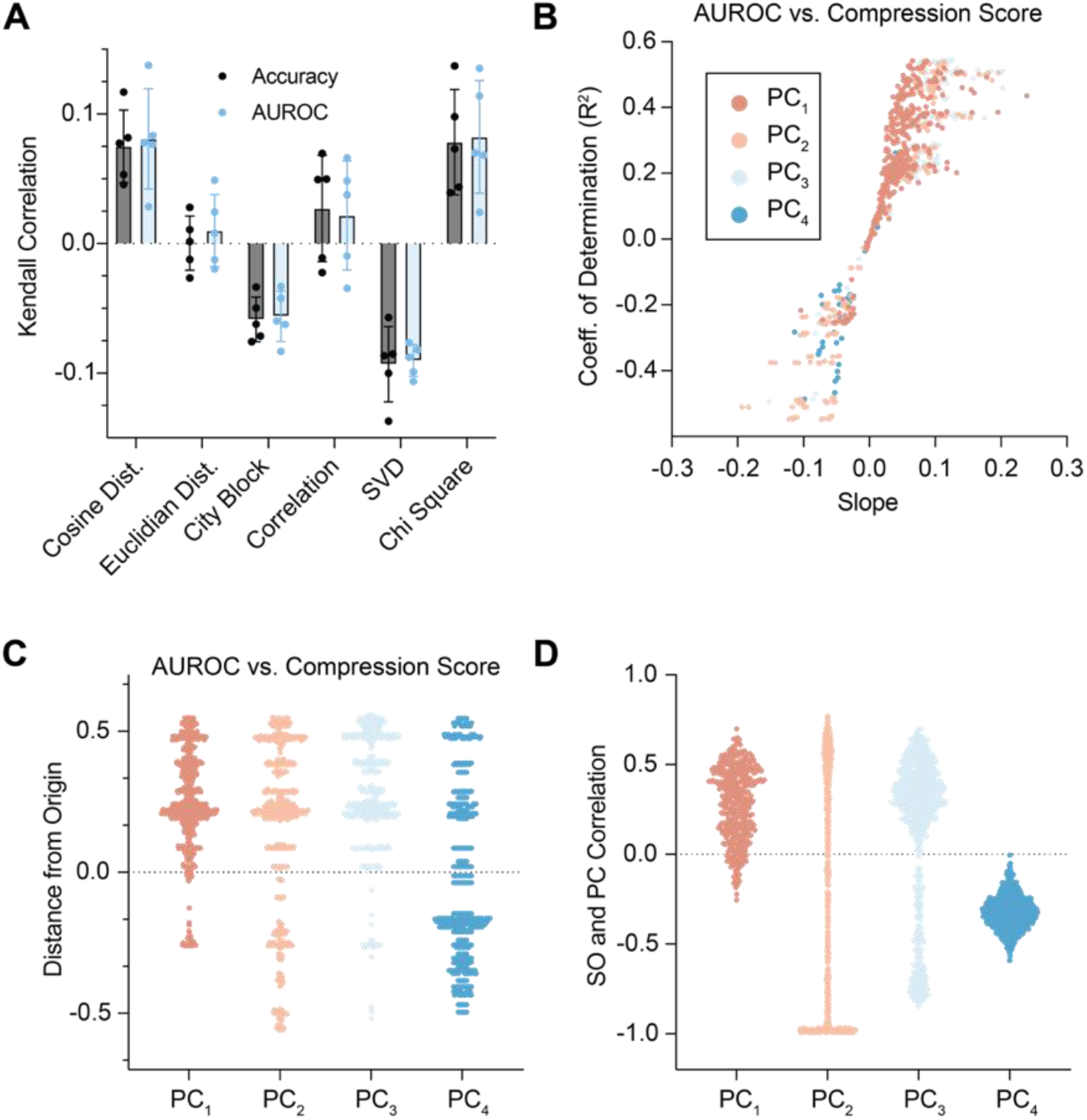
Comparing various metrics for substrate orthogonality and protease coverage. **(A)** Scoring various metrics that quantify substrate orthogonality, where the metrics are listed on the x-axis. Kendall correlation (y-axis) is measured for simulated libraries (n = 100) between the substrate orthogonality metric (x-axis) and classification metric (AUROC or accuracy). Classification challenge is simulated based on microarray gene expression data of a healthy and disease murine sample. This challenge is repeated 5 times for each metric. Cosine distance is selected for use in the compression score. **(B)** Comparing 4 metrics of protease coverage (i.e., PC_1_, PC_2_, PC3, PC4). Each dot represents one scoring run, in which 290 simulated libraries, each comprising 50 substrates are tested in a simulated classification challenge. For all tested libraries, a coefficient of determination (R^2^) and slope are calculated for the plot of AUROC vs. Compression score metric, which uses one of four protease coverage (PC) metrics. Points with high R^2^ and slope indicate that the protease coverage metric is predictive of classification performance. **(C)** Quantifying distance from origin (−0.3, −0.5) for all points in panel B. Each protease coverage metric is plotted on the x-axis and distance is plotted on y-axis. Higher distance values indicate the protease coverage metric is predictive of classification performance. **(D)** Quantifying correlation (Pearson’s *r*) between each protease coverage metric and the substrate orthogonality metric (i.e., cosine distance). Smaller correlation values indicate that the protease coverage metric is providing unique information from the substrate orthogonality metric.

**Figure S3.**
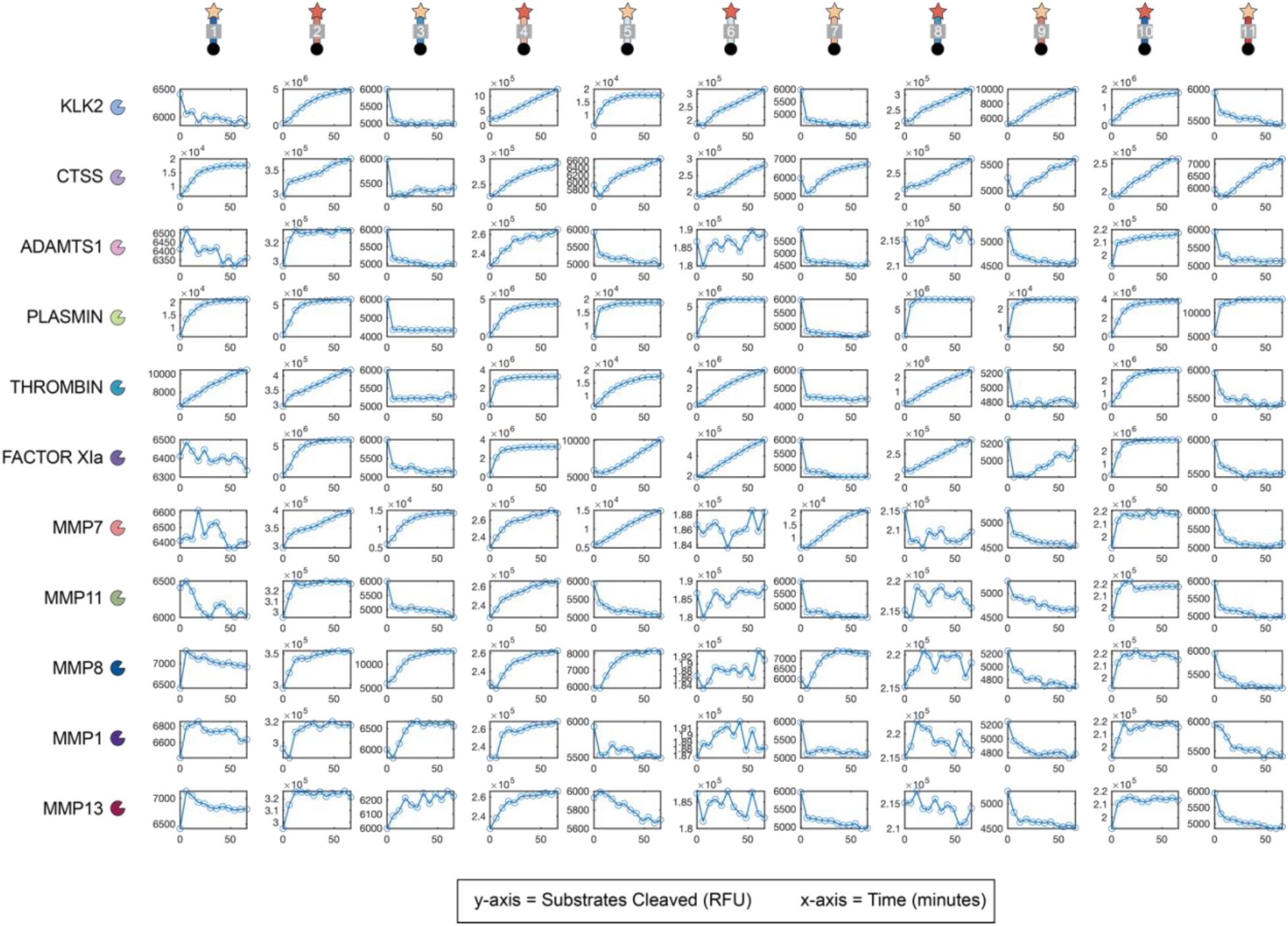
Raw kinetic substrate cleavage data for all protease-substrate combinations. Each plot is the raw fluorescence data from a protease cleavage assay between a particular protease (row) and substrate (column). Each plot shows the amount of substrates cleaved, as quantified by fluorescence (RFU), on the y-axis, over time (x-axis; minutes). Maximum velocity, *V_max_*, or product formation rate (RFU/min) is extracted and plotted as single value in heatmap in the main text.

**Figure S4.**
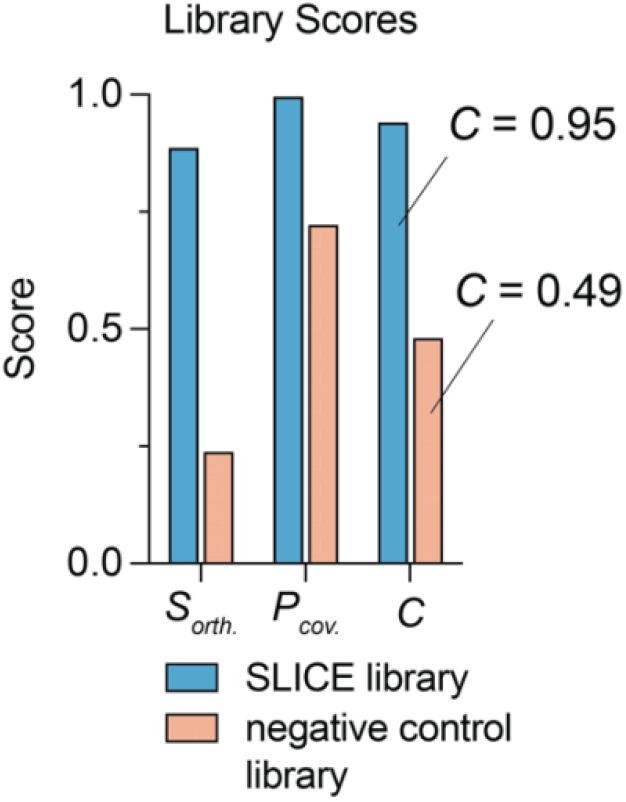
Comparing library scores for SLICE and negative control. The normalized score value (y-axis) for substrate orthogonality (*S_orth_*), protease coverage (*P_cov_*), and compression (*C*), are compared for a SLICE library (blue bars) and the negative control library (orange bars). The SLICE library has an overall compression score of C = 0.95, whereas the negative control library has a compression score of C = 0.49.

**Figure S5.**
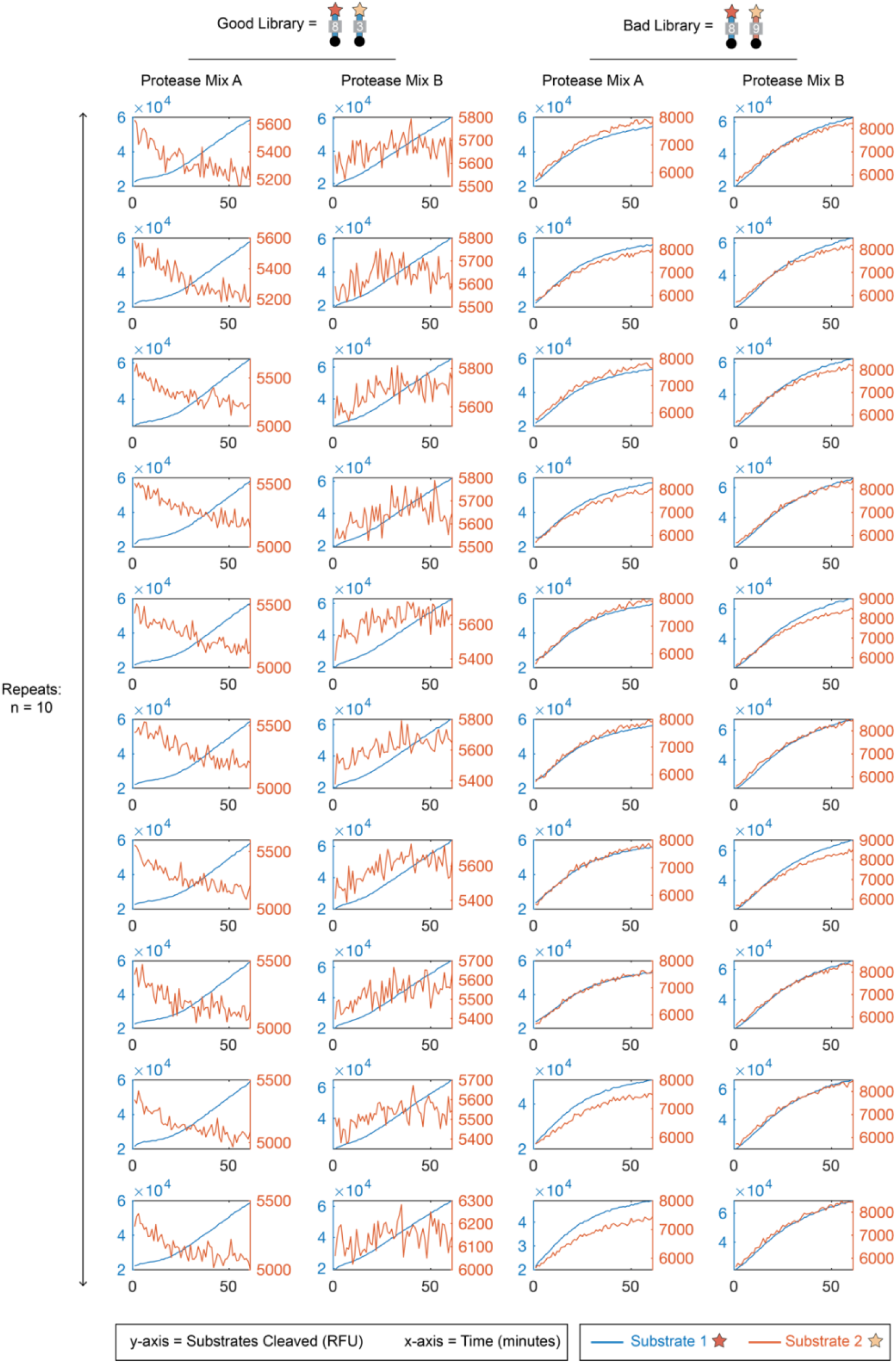
Raw kinetic substrate cleavage data from *in vitro* classification challenge. Each plot represents the fluorescent read-out of substrate cleavage from one well from the *in vitro* classification challenge. Each well included (1) a mixture of 11 proteases according to a concentration profile (i.e., Protease Mix A, Protease Mix B) and (2) a library comprising 2 fluorogenic peptide substrates, as indicated by the column. The same conditions for each column were repeated 10 times each (rows). Each plot shows the amount of substrates cleaved, as quantified by fluorescence (RFU), on the y-axis, over time (x-axis; minutes). Maximum velocity, *V_max_*, or product formation rate (RFU/min) is extracted and plotted as single value for each substrate in the main text. In each plot, the signal from Substrate 1 (i.e., 5-FAM substrate) is plotted as the blue trace (left y-axis), and the signal from Substrate 2 (i.e., EDANS substrate) is plotted as the orange trace (right y-axis).

**Figure S6.**
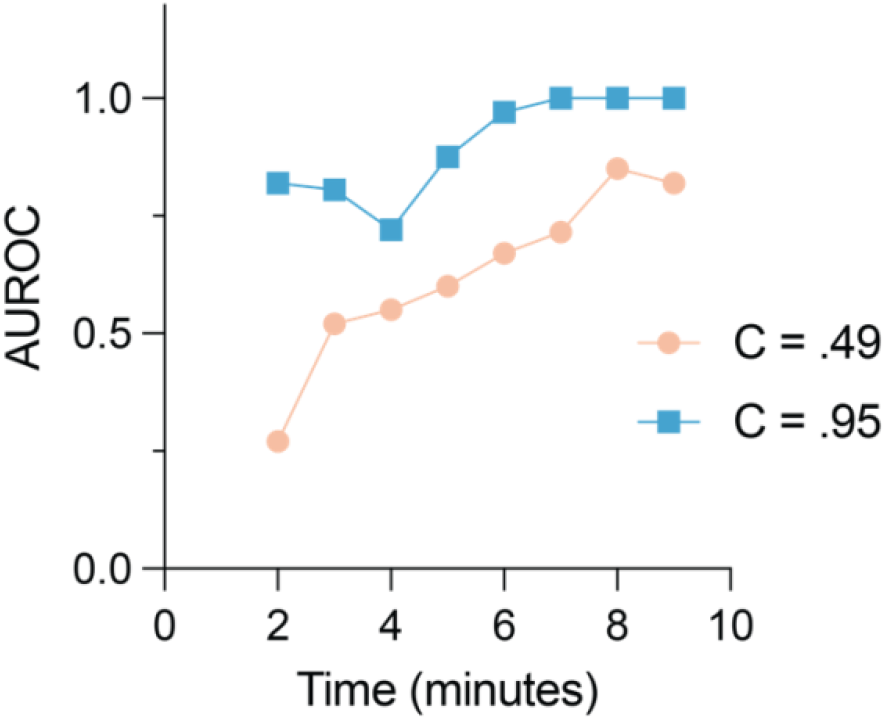
Comparing classification power at various end points. Each point represents the resulting classification performance (y-axis; AUROC) of a substrate library where the product formation rates for each substrate signal is calculated with a different end-point in time (x-axis; minutes). The SLICE library (C = .95) is plotted as the blue trace with square markers and the negative control library (C = .49) is plotted as the orange trace with circle markers.

**Figure S7.**
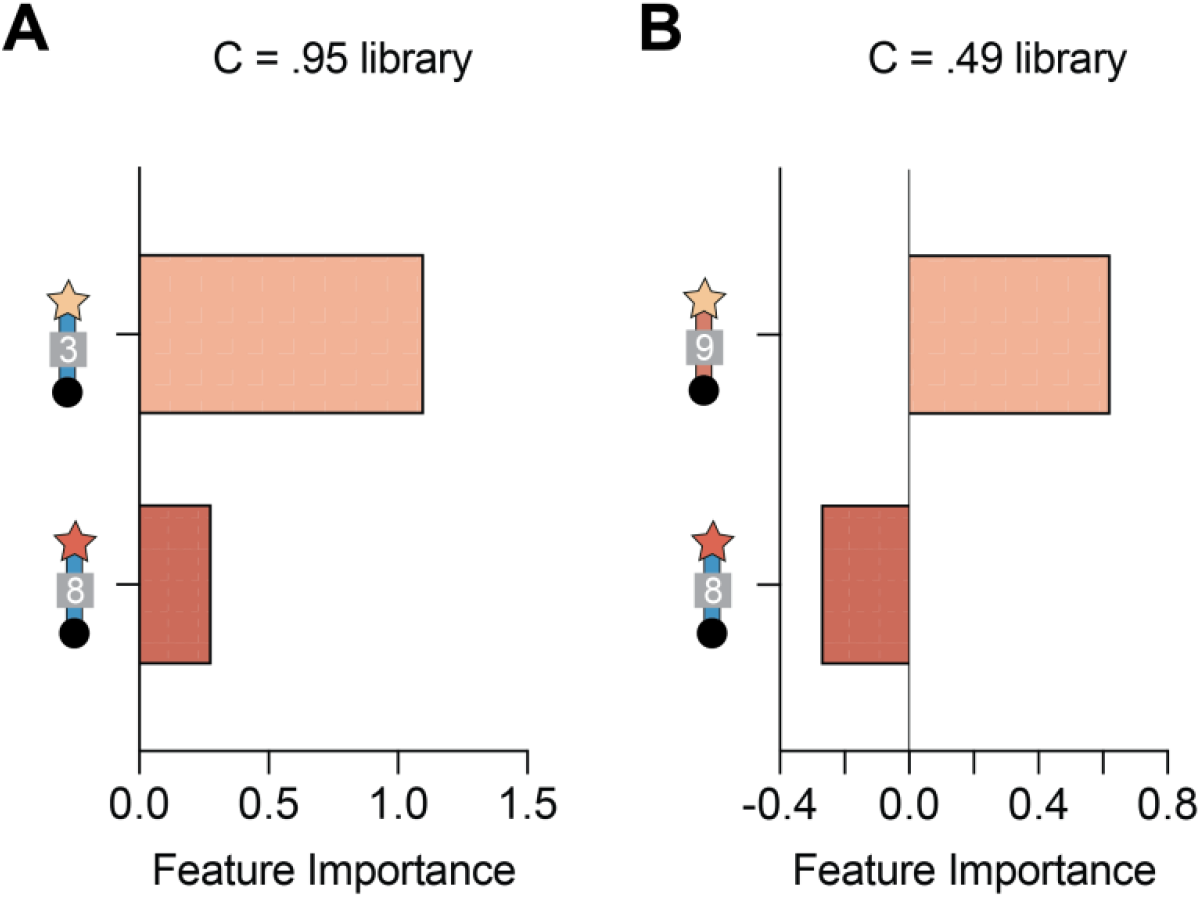
Comparing feature importance score for each substrate signal. Plot of the feature importance scores for each substrate signal in the **(A)** C = .95 library and the **(B)** C = .49 library. The light orange bar represents the EDANS substrate, and the dark red bar represents the 5-FAM substrate. Feature importance score is extracted from training the random forest model in classifying mixture A from mixture B in the *in vitro* classification challenge.

